# CT-Derived *In Vitro* Model Simulating Neonatal Cranial Growth

**DOI:** 10.64898/2026.11.17.767179

**Authors:** Ahmet Arif Colakoglu, Yunus Samiepour Ardakani, Sina Najjari, Ce Liang, Anastasiia Maliuk, Roman Hossein Khonsari, Hans Delye, Mehran Moazen, Peter Culmer, Ali Alazmani

## Abstract

**Purpose:** This study presents an *in vitro* platform for quantitatively simulating cranial growth in neonates from 3 to 6 months of age.

**Methods:** The system integrates a pneumatic–hydraulic actuation unit, multi-camera optical tracking, and a CT-compatible enclosure to replicate and measure intracranial expansion. A phantom head model was fabricated via 3D printing using cranial bones and flexible latex sutures, reproducing neonatal cranial biomechanics. Controlled hydraulic pressure simulates intracranial volume (ICV) growth, while surface and volumetric changes are captured through synchronized imaging.

**Results:** Morphometric parameters – biparietal diameter, occipitofrontal diameter, cranial circumference, and ICV – were measured and compared against published reference data. The system achieved high reproducibility across repeated trials. CT-derived ICV values (***R***^2^ **= 0.98**) closely matched reference trends, with average manufacturing deviations of 0–4 mm between fabricated and reference geometries. Linear morphometric increments confirmed physiologically consistent expansion patterns.

**Conclusion:** The proposed platform provides a quantitatively validated, controllable, and CT-compatible environment for studying neonatal cranial development. It enables biomechanical testing and device evaluation under ethical and repeatable *in vitro* conditions, with potential future applications in surgical training and pre-operative planning.

## 1 Introduction

Cranial growth in healthy neonates is a vital developmental process, primarily driven by rapid brain expansion and governed by the biomechanical interplay between cranial bony plates and cranial joints or sutures [1–4]. During the first year of life, the brain undergoes rapid enlargement, doubling in volume from approximately 400 ml at birth to nearly 800 ml by six months, necessitating corresponding skull expansion to accommodate this growth [5]. This early period is especially critical, as proper cranial development supports healthy neurological function [6]. A clear understanding of these dynamics is essential for diagnosing and managing clinical conditions such as craniosynostosis, where premature suture fusion impedes growth [7–9], and epilepsy, where cranial protection during seizures is a concern [10]. Such knowledge informs clinical strategies for monitoring skull development and designing intervention tools.

Research into cranial growth spans human infant studies, computational in silico models, and animal models, each offering unique strengths and limitations [1, 2, 11–13]. Human studies provide high anatomical accuracy and direct clinical relevance, offering valuable insight into natural growth patterns [5]. However, they are constrained by ethical restrictions, small sample sizes, significant costs, time-intensive data collection, and inter-individual variability [14, 15]. In silico models offer a cost-effective and ethically unrestricted means to simulate cranial development with parametric control. Yet, they often oversimplify key biomechanical factors such as suture flexibility and intracranial pressure dynamics and require careful validation [16, 17]. Animal models — principally rodents (mice and rats), rabbits, and pigs — permit controlled, longitudinal experimentation and have advanced understanding of suture biology and calvarial bone formation [11, 13, 18]. However, their anatomy differs substantially from the human neonatal skull: rodents are small and lissencephalic with a quadrupedal skull-base orientation; rabbits share these quadrupedal proportions; and the pig, although closer in overall size, has a snout-dominated craniofacial form and a thicker, structurally distinct cranial bone architecture that makes it a poor geometric analogue of the human calvaria. These anatomical differences, together with ethical and financial constraints, limit the direct translation of animal data to human cranial development [18]. Moreover, while human studies are limited by inter-individual variability, existing computational and animal approaches remain largely static, offering no controlled and repeatable means of reproducing the dynamic expansion of the neonatal skull [19].

Current technologies designed to manage cranial growth such as medical devices and wearable orthotic helmets are used to guide skull morphology in conditions like craniosynostosis or to offer cranial protection in cases such as epilepsy [20, 21]. Several medical device companies have developed orthotic helmets specifically for the treatment of craniosynostosis, applying gentle external forces to direct cranial expansion during early infancy [22–28]. These interventions require robust testing platforms to evaluate safety, efficacy, and fit. However, testing such devices on human subjects is restricted by ethical and practical constraints, and conventional experimental setups often fail to replicate realistic cranial biomechanics [29–32]. This limitation also affects hands-on surgical training, where the absence of anatomically accurate, dynamically growing skull models hinders the development and evaluation of patient-specific clinical strategies [33–37].

To address these challenges, this study introduces a dynamic *in vitro* model designed to simulate the cranial growth of a healthy neonatal skull from 3 to 6 months of age. Unlike static models, this system replicates the biomechanical properties of the developing skull, combining calvarial bone plates with flexible sutures to enable anatomically realistic expansion. It offers a controlled, repeatable, and ethically sound platform for biomechanical research and the testing of therapeutic devices such as cranial orthoses, with broader potential applications in surgical training and education [38]. Growth targets are based on age-specific reference intracranial volumes (ICV) from a normative craniofacial growth model — for example, a reference ICV of 743.13 ml at 4 months of age [1] — ensuring developmental fidelity. This paper outlines the design and fabrication of the model, along with the experimental methodology and performance evaluation, with the aim of establishing a novel, anatomy-derived and potentially patient-adaptable platform for simulating early cranial development (Fig. 1).

**Fig. 1.**
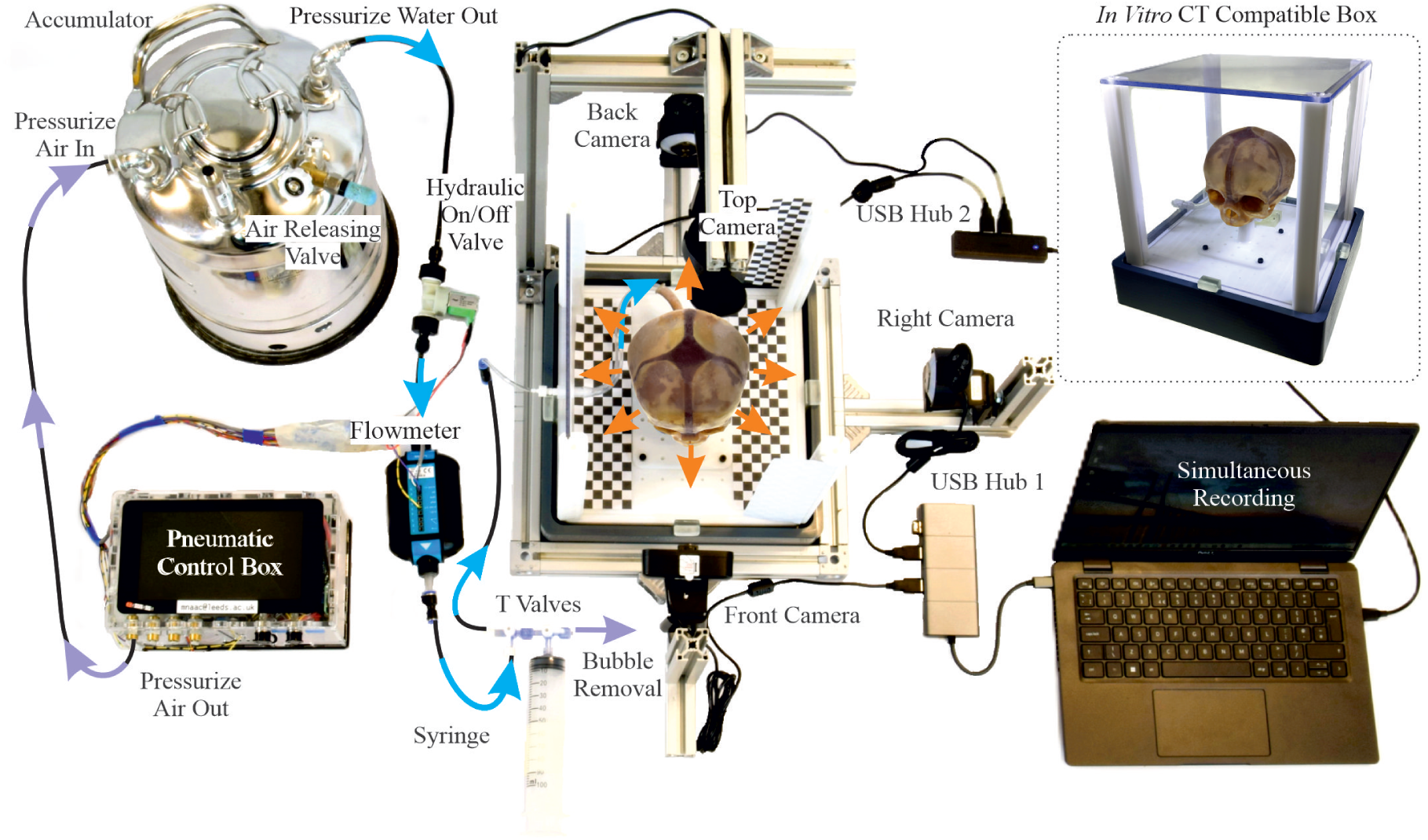
Overview of the *in vitro* actuation system used to simulate neonatal cranial growth. The setup consists of a pneumatic–hydraulic control box with GUI, connected to a 10-L accumulator for pressure-driven water delivery through a hydraulic valve and flowmeter. A syringe and T-junction valves enable bubble removal. Tubing interfaces with the head model through the foramen magnum, inflating the internal latex chamber to induce expansion. The model is housed in a CT-compatible enclosure and monitored by four cameras (superior, lateral, posterior, anterior).

## 2 Materials and Methods

An *in vitro* experimental setup was developed to simulate the physical growth of a healthy neonatal skull from 3 to 6 months of age, a period of rapid cranial expansion starting from the available reference segmentation [1, 5]. This setup emphasizes biomechanical accuracy by selecting materials that approximate the elastic modulus and other mechanical properties of infant skull and soft tissues, while allowing controlled dynamic expansion. It integrates three subsystems: a 3D printed model of an individual head, an actuation system for pressure-driven expansion, and a multi-modal measurement system for data acquisition. To ensure fidelity, several design requirements were established. The head model was required to replicate neonatal geometry using rigid materials for cranial bones, flexible latex for sutures and skin layers, and dissolvable supports for assembly. The actuation system needed to deliver precise fluid flow and pressure to achieve age-specific ICV increments, using the 3- to 6-month reference range as target stop points, with mechanisms for flow regulation and bubble removal. The measurement subsystem had to support surface-deformation tracking via multi-angle cameras and volumetric analysis via computed tomography (CT), while maintaining CT compatibility to avoid artefacts, preserve image quality, and prevent equipment damage.

To align with clinical practice, where surgeons routinely measure distances and cranial circumferences manually, a tri-modal measurement approach was implemented [39, 40]. Linear distances and cranial circumferences were first measured manually using calipers and tape. Optical tracking with multi-angle cameras was used to monitor surface deformation, followed by CT imaging for volumetric assessment. This combination provided cross-validated data and ensured repeatability within an ethical *in vitro* framework.

### 2.1 Physical Head Model and Actuation System

Age-specific cranial geometries for neonates at 3, 4, 5, and 6 months were obtained from Liang et al. [1, 5]. These digital reference models are derived from clinical CT scans of 217 normally developing individuals [5] and have been biomechanically validated through a physico-mechanical finite element model of postnatal craniofacial growth that uses ICV-driven calvarial expansion as the primary growth driver, achieving surface deviations within ±5 mm for over 80% of the skull surface [1]. The 3-month geometry served as the baseline for fabricating a single physical phantom skull, while the 4- to 6-month geometries provided independent reference targets for evaluating phantom expansion. Two modifications were applied to the baseline geometry to improve functionality: (1) fixation points were integrated at the basion to secure the skull during experiments with minimal influence on expansion, and (2) 2 mm holes were added across cranial bone surfaces to allow integration of the inner and outer latex layers, enhancing uniform expansion (Fig. 2).

**Fig. 2.**
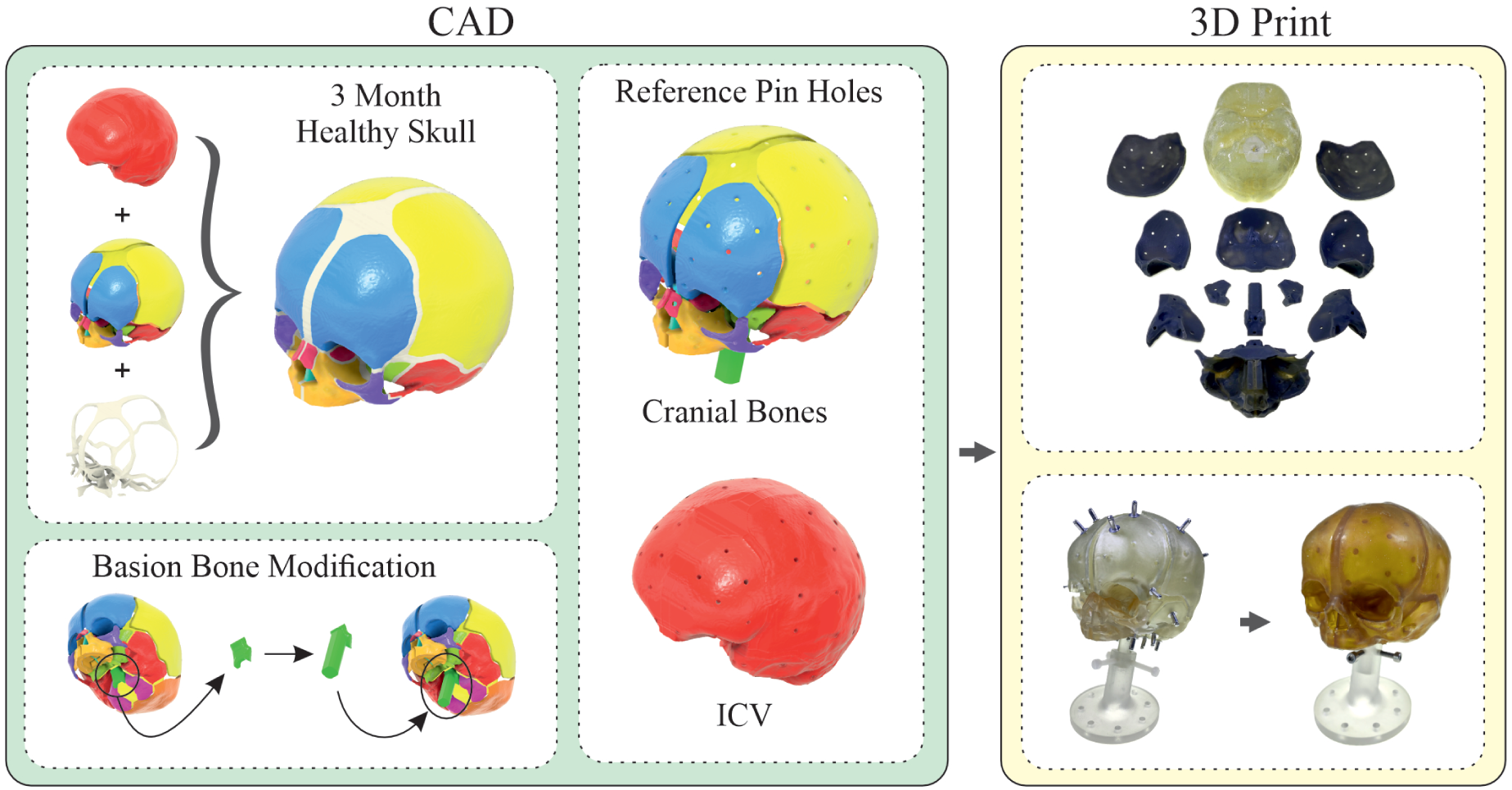
Rendering from Autodesk Fusion 3D showing key modifications to the 3-month skull model. A fixation feature was added at the basion to stabilise the skull during growth with minimal impact on expansion. Multiple 2-mm holes were incorporated to enable integration between the inner and outer latex layers of the head model.

The cranial bones were fabricated via stereolithography (SLA) (Formlabs Clear Resin V4) to approximate neonatal bone stiffness [41, 42]. The ICV was printed via fused deposition modelling (FDM) using dissolvable polyvinyl alcohol (PVA) on a 3D printer (X1-Carbon, Bambu Lab) to serve as a temporary scaffold. Bones were aligned around the scaffold using matching alignment holes to maintain anatomical accuracy. Latex was first applied externally to form sutures and skin, filling the holes for integration. After curing, the PVA scaffold was dissolved in water, and a second latex layer was applied internally, bonding through the perforations to form a sealed expansion chamber. Finally, latex tubing was integrated at the foramen magnum to provide a realistic fluid-filled interface for actuation. The materials selected for the physical head model fall within the reported biological property ranges of human head tissues, and the corresponding validation data, along with initial unsuccessful manufacturing attempts, are provided in in Appendix B and Appendix C.

The actuation subsystem was designed as a multifunctional pneumatic control box capable of precise pressure and volume regulation [43]. (Full details in Appendix A.) It incorporates a Raspberry Pi 4B microcontroller with custom firmware and a touch-screen graphical user interface (GUI) for real-time adjustment of pressure and flow parameters. Four pneumatic output channels (0–150 kPa) provide regulated pressure to drive water from a 10-L accumulator through high-pressure biocompatible tubing into the phantom. Solenoid valves (V114A-5MUB, SMC, Japan) and pressure transducers (TruStability SSC-D-AN-N-015PG-2-A-3, Honeywell), mounted on each channel via T-connectors, enable accurate control and monitoring, while an automatic hydraulic on/off valve prevents over-expansion by closing at the programmed volume [44]. A digital flowmeter (Atrato 710, Titan Enterprises) tracks fluid delivery to meet age-specific ICV targets. A syringe and T-junction valves allow vacuum-assisted bubble removal. Water was selected as the actuation medium due to its incompressibility and consistent volume delivery [45–48]. USB ports allow data logging and integration with external sensors. A detailed schematic is provided in Appendix A (Fig. 7).

### 2.2 Measurements and Experimental Design

A multi-modal measurement approach was used for the detailed assessment of cranial growth, integrating manual cranial morphometric measurements, multi-camera optical measurement, and CT imaging. Manual anthropometric measurements – including cranial circumference, biparietal diameter (BPD), and occipitofrontal diameter (OFD) – were performed using standard clinical tools, following established anatomical landmark protocols (the glabella and opisthocranion for OFD and the parietal eminences for BPD) [39, 40, 49–54]. All manual measurements were carried out by a single operator (linear dimensions with calipers and cranial circumference with tape) to remove inter-observer variability, using the same anatomical landmarks throughout to maintain consistency across ages. At each age/ICV condition, every dimension was measured three times, and all values are reported as mean ± standard deviation (*n* = 3; Appendix D, Table 3), with a coefficient of variation below 1.7% for all manual measurements and below 1.0% for all optical measurements. Four webcams were positioned around the phantom to capture superior, lateral, posterior, and anterior views for real-time observational monitoring and multi-camera optical measurement of surface deformation, complementing the CT-based volumetric measurements which served as the primary three-dimensional reference. Each camera was treated as an independent two-dimensional view; a stereophotogrammetric setup with overlapping fields of view and a common extrinsic calibration was not employed. Because each view records a three-dimensionally expanding surface as a two-dimensional projection, these measurements are subject to out-of-plane motion, perspective distortion, and parallax error. Consequently, only a pixel-to-millimetre scale factor was determined for each view, using the CT-derived BPD of the 3-month model for the superior, posterior, and anterior views and its CT-derived OFD for the lateral view, and no three-dimensional surface reconstruction was attempted from the camera data. These two dimensions are therefore identical to their CT values at the 3-month baseline (Table 3), and independent agreement between the modalities is assessed from 4 months onward. This was a deliberate design choice: CT imaging provided the primary three-dimensional ground truth, so the camera array was intended for real-time monitoring and supplementary in-plane surface measurement rather than for volumetric reconstruction.

A CT-compatible enclosure (30 cm × 30 cm × 30 cm) housed the phantom head, allowing direct placement into a CT scanner (XT H 225 ST, Nikon, Japan) for volumetric analysis. CT scans were performed using 150 kV, 150 *µ*A, a 50 *µ*m voxel size, 2000 projections, and a 30-minute scan time; including phantom positioning, each acquisition required approximately 50 minutes. During CT trials, iodine solution replaced water to improve ICV segmentation contrast. CT data were processed (Avizo 2022.1, Thermo Fisher Scientific, USA) [55] to extract ICV, generate sectional views, reconstruct 3D geometries, calculate cephalic index (CI), and produce colormaps illustrating shape changes and manufacturing deviations.

Two experimental protocols were performed: one camera-based and one CT-based. Both used sequential actuation to reach target ICVs at 3, 4, 5, and 6 months [1, 5]. For camera experiments, tubing was connected to the foramen magnum and water was injected to approximately 720 mL. Bubbles were removed by vacuum before adjusting the volume to the precise 3-month target (694.93 mL). Pressure was then regulated via the GUI to reach each subsequent ICV target (743.13, 788.29, and 830.51 mL). Cameras recorded deformation at each stage, and videos were analyzed in video analysis software (Kinovea, 2023.1.2 version) [56]. In Kinovea, tracked point displacements are reported in pixels and require a spatial scale reference to be converted into metric units. It would be inappropriate to use the background checkerboard for this purpose because it lies on a different plane and depth from the skull. Instead, the pixel-to-millimetre scale for each camera view was defined using known dimensions of the 3-month model measured from its CT reconstruction in Avizo, in which the reference corresponds to the object plane of the skull itself and the scale is relevant to the measured surface. Cranial circumference was also extracted from the camera footage but was reported from the manual measurement, because planar 2D views of a 3D object are inherently subject to errors (i.e. due to perspective and parallax effects). As internal pressure increased, the skull underwent both lateral widening and a vertical upward displacement; this displacement shifted the model relative to the pre-defined camera angles and scale plane, introducing out-of-plane motion, perspective distortion, and parallax error, and intermittently obscuring the landmarks required to trace the full circumference. For CT experiments, the same preparation and actuation procedure was used, with iodine solution for improved image contrast. The enclosure was scanned at each ICV increment, and segmented data were used to obtain volumetric and shape-based metrics. All experiments were repeated to ensure consistency and cross-validation across modalities.

## 3 Results

### 3.1 Head Model and Actuation System

The assembled head model using 3D printed bones and covering both inner and outer surfaces with latex layers was tested by air and water, respectively in terms of flexibility and leak-proof before conducting experiments in the *in vitro* setup environment.

The actuation subsystem successfully simulated intracranial expansion in the phantom head model, accurately replicating neonatal cranial growth dynamics. An example of the full expansion is provided in supplementary (Online Resource Video 1.) Fig. 1 shows the complete setup, including the electro-pneumatic regulator (ITV1010-21F2BL5, SMC, Japan), 10-L accumulator, digital flowmeter, 100-mL precision syringe, camera system, computer, and skull model. It also illustrates the CT-compatible box housing the phantom for imaging.

The electro-pneumatic regulator maintained pressure stability within ±0.1 kPa across the 0–150 kPa range, verified using the integrated pressure transducers (TruStability SSC-D-AN-N-015PG-2-A-3, Honeywell) [57], which were connected to each pneumatic channel via T-connectors as described in Appendix A. A single pneumatic channel pressurized the accumulator, delivering water through tubing to the phantom’s foramen magnum. Solenoid valves controlled flow via PWM signals, while the hydraulic on/off valve (0.05-mL resolution) prevented over-expansion. The flowmeter, calibrated for 0.1–10 mL/s, reliably achieved age-specific ICV targets of 694 mL (3 months) and 743 mL (4 months) [1, 5]. Air bubble removal using a syringe-driven vacuum cycle ensured stable fluid transmission. The GUI on a 10-inch touchscreen (1280 × 800) allowed real-time control of pressure, experiment duration, and flow graphs. Data were logged directly to the microcontroller and exported via USB. Three-dimensional analysis in Fig. 3 shows manufacturing error by age across five anatomical views. Comparing the phantom geometry with the reference model revealed that the 3-month model aligned closely with the reference (predominantly blue), while the 6-month model showed increasing red/orange regions, indicating approximately 3-mm geometric deviation primarily in the parietal region.

**Fig. 3.**
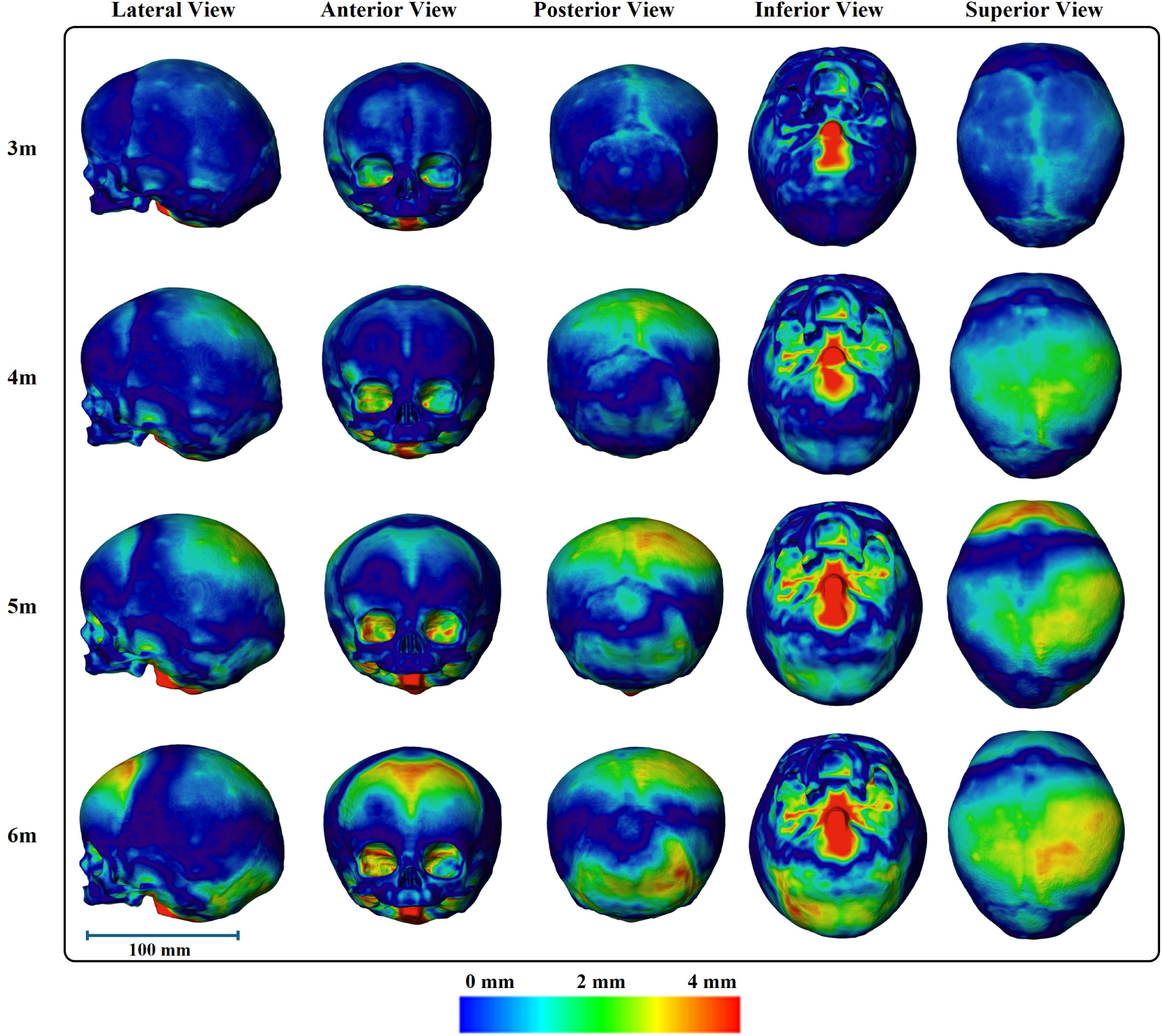
Colormap showing deviations between the phantom and reference skulls (3–6 months) across five standardized views. Colors represent errors from 0 mm (blue) to 4 mm (red). All models are shown at the same scale: a 100 mm scale bar is provided for reference.

### 3.2 Cranial Morphometric Measurements

Manual anthropometry provided direct measurements of external cranial dimensions across superior, lateral, posterior, and anterior projections. Key parameters included BPD, OFD, cranial heights (LCH, PCH, ACH), cranial circumference, and CI. As summarized in Appendix D (Table 3), these values showed consistent age-related increases. For example, BPD increased from 113.35 ± 0.86 mm at 3 months to 120.32 ± 0.85 mm at 6 months, reflecting transverse expansion, while OFD increased from 137.13 ± 1.81 mm to 143.09 ± 1.27 mm, indicating anteroposterior elongation. Cranial circumference increased from 400.33 ± 2.52 mm to 416.67 ± 4.04 mm, and CI ranged from 82.7–84.1%, indicating mild brachycephalic stabilization in early postnatal stages.

Multi-camera optical measurement provided supplementary in-plane measurements of surface deformation from four independent views. Fig. 4 shows representative images from the four camera views across all ages, with green lines marking transverse widths and orange lines marking cranial heights. Multi-camera optical measurement results showed (Appendix D, Table 3) progressive increases in cranial widths (e.g., BPD: 113.14 ± 0.00 mm at 3 months, the calibration reference, to 124.31 ± 0.56 mm at 6 months) and cranial heights (e.g., PCH: 112.03 ± 0.26 mm to 121.87 ± 0.82 mm). This modality captured directional growth patterns, with transverse expansion occurring slightly more rapidly than height increases. Occipitofrontal diameter agreed closely with manual measurements (within 1.0% at all ages), whereas anterior cranial height exceeded the manual value by 11.5–23.1% and cranial length measured from the superior view was consistently shorter than from the lateral view; both discrepancies reflect view-dependent landmark visibility and perspective distortion.

**Fig. 4.**
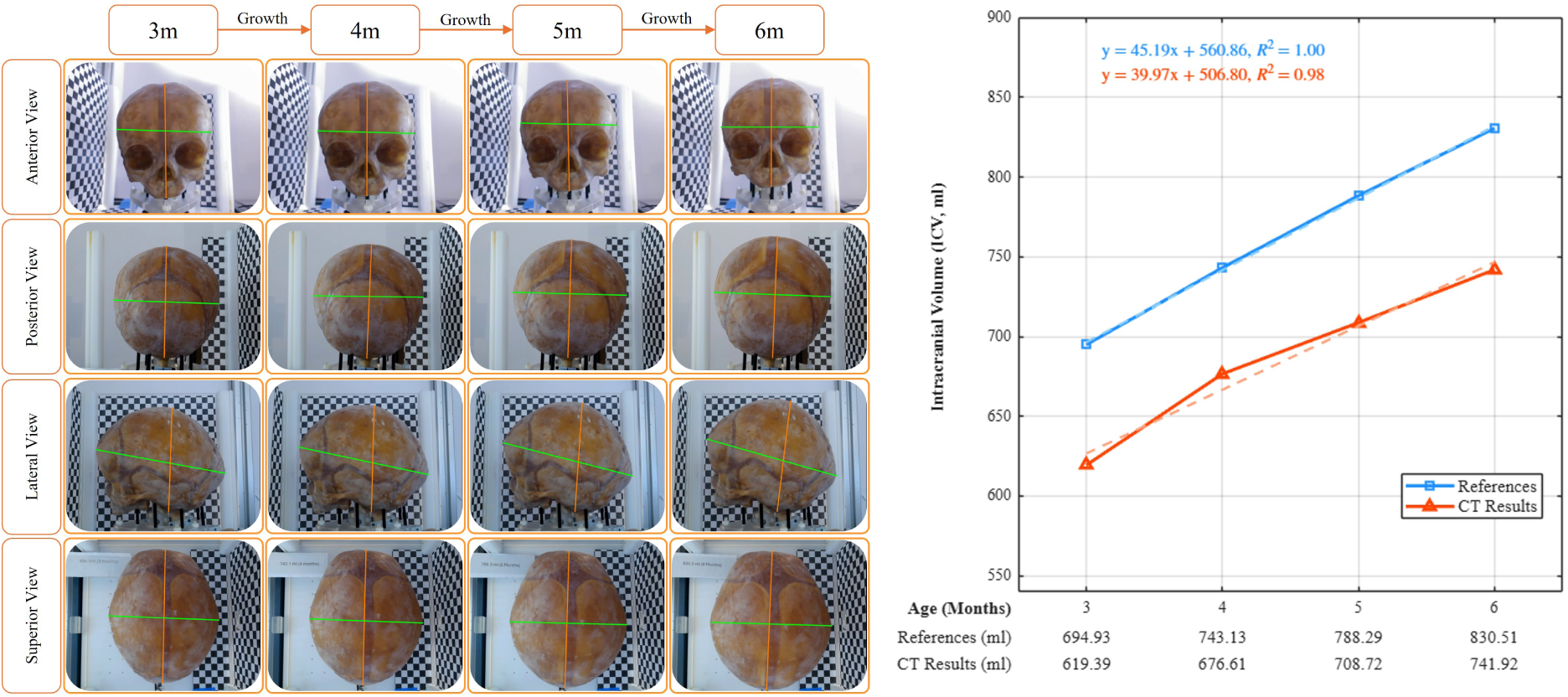
Cranial growth from 3 to 6 months based on camera recording. Four views (anterior, posterior, lateral, superior) are shown vertically, with age progression displayed horizontally. Cameras were calibrated using CT dimensions of the 3-month model, and subsequent stages were tracked using the software tool. Green lines indicate cranial widths and orange lines indicate cranial heights. The accompanying plot shows intracranial volume (ICV) for reference data (blue) and CT-derived results (orange) from 3–6 months, with linear trend lines and equations for both datasets.

CT-derived ICV measurements provided volumetric insight into cranial capacity. Fig. 4 plots ICV values for reference and experimental data with linear regression fits. Reference ICV increased from 694.93 mL at 3 months to 830.51 mL at 6 months, whereas the CT-derived ICV of the phantom increased from 619.39 mL to 741.92 mL over the same range, remaining consistently below the reference targets by approximately 75–90 mL. This offset is not a discrepancy between imaging modalities or between separate models – a single physical phantom was used throughout – but arises primarily from the fabrication of the phantom: its internal cavity is lined with a manually cast latex layer (the brain-shaped balloon) that acts as a dura-like inner membrane and occupies part of the internal space, so the resulting fluid-filled cavity is slightly smaller than the digital target geometry. A smaller, secondary contribution comes from segmentation variability, as the iodine-contrast threshold used to delineate the cavity in Avizo affects the absolute extracted volume. Nevertheless, both datasets closely follow parallel linear trends, consistent with the expected encephalic-driven cranial expansion of early infancy, so this systematic offset does not alter the growth trajectory reproduced by the phantom [40]. All measurements reported here were obtained from this single phantom; the variability they reflect is therefore the consistency of repeated experiments on the same phantom rather than variation across different models.

### 3.3 Comparative Evaluation and Visual Analysis

Appendix D (Table 3) consolidates morphometric measurements across all modalities, allowing direct comparison across methods. BPD values show close agreement between manual anthropometry and CT (e.g., at 5 months: 118.71 ± 1.95 mm vs. 118.58 mm), while multi-camera optical measurement is slightly higher (120.77 ± 0.29 mm). This difference increases with age and is consistent with out-of-plane motion and parallax error. OFD values are highly consistent across modalities, with differences below 1% (e.g., 141.65 ± 1.47 mm manual vs. 140.84 ± 0.67 mm multi-camera optical measurement at 5 months), indicating stable anteroposterior measurements. PCH and ACH are also comparable, with CT values generally slightly higher. This small difference may arise from the challenge of positioning manual reference points identically for each measurement, reflecting a natural variation between methods. Global measures such as cranial circumference (e.g., 414.00 ± 5.00 mm manual at 5 months) and CI (81.8–87.7% in multi-camera optical measurement) showed progressive cephalic broadening, with typical age-related increments of 2–7 mm and 1–3%, respectively. High correlations between references and CT measurements (e.g., *R*^2^ = 0.98 from Fig. 4, comparative plot for CT results) demonstrate high consistency across modalities and supports the reliability of the collected dataset.

Fig. 5 displays monthly growth relative to previous ages, with increasing yellow, orange, and red areas representing up to 4-mm outward expansion by 6 months. These visualizations collectively demonstrate controlled and repeatable cranial vault expansion that approximates the physiological growth pattern targeted in this study, arising from the prescribed volumetric actuation together with the elastic deformation of the phantom rather than from osteogenesis [40].

**Fig. 5.**
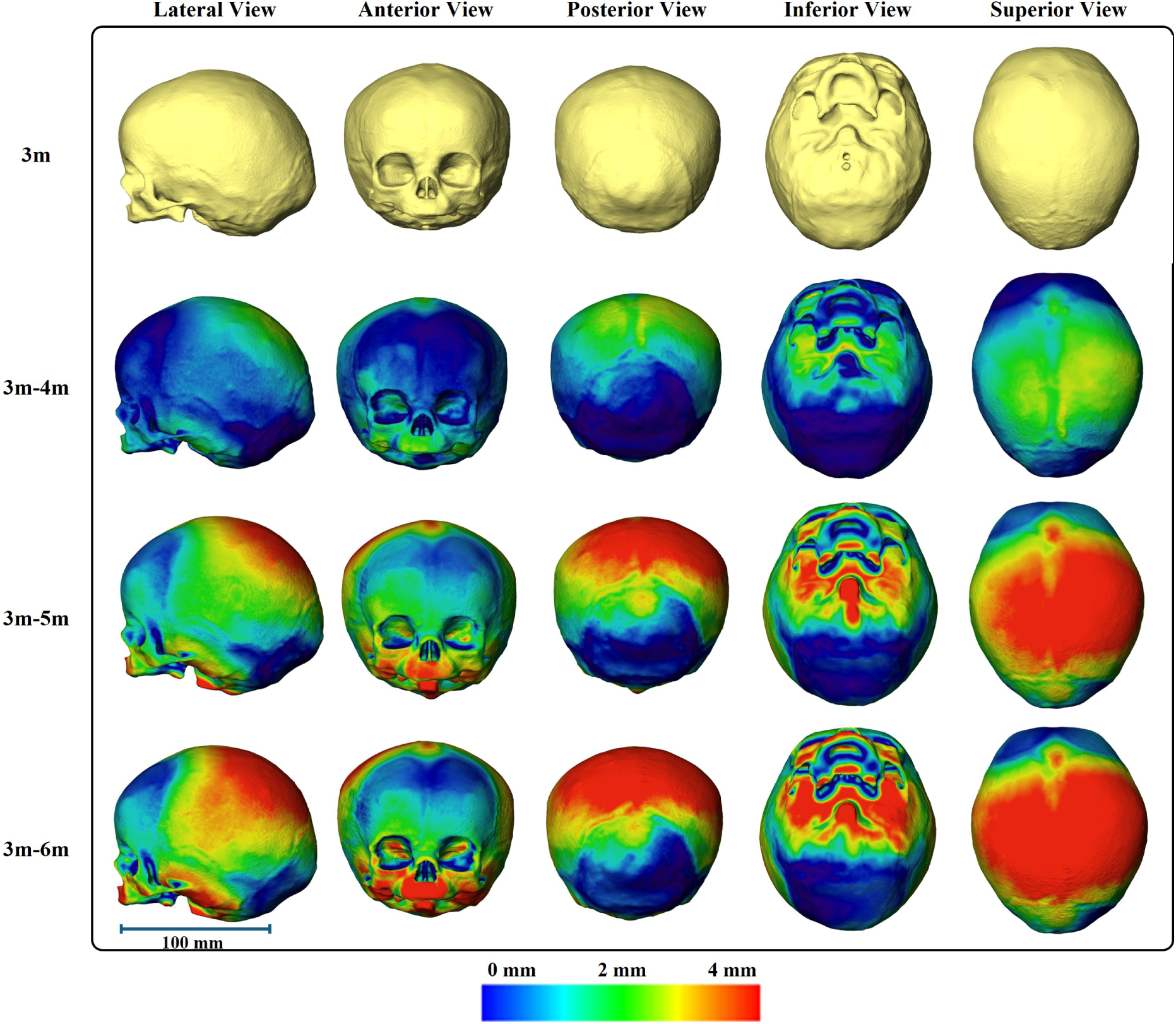
Colormaps showing monthly skull growth (3–6 months) across five anatomical views. Deviations from the previous age are mapped from blue (0 mm) to red (4 mm). All skulls are shown at the same scale with a 100 mm reference bar. Increased yellow/orange/red areas, especially in the parietal region, indicate up to 4 mm outward expansion relative to the 3-month model.

Two-dimensional sectional CT views enabled detailed geometric comparison with reference models [1, 5], as illustrated in Fig. 6. CT results across 3–6 months showed expected expansion patterns, while reference 3–4 month growth provided a baseline for comparison. Sagittal (P1) views demonstrated that CT reconstructions showed slightly greater anterior–posterior elongation than reference models. Coronal planes (P2, P3) highlighted minor height increases; P2 maintained bilateral symmetry with *<*1 mm deviations, while P3 showed 2–3 mm elevations in the posterior region. Horizontal (P4) sections confirmed circumferential uniformity, with subtle asymmetries more visible in CT than in reference geometries. Collectively, the planes illustrate anisotropic growth, with sagittal and coronal expansion exceeding horizontal widening.

**Fig. 6.**
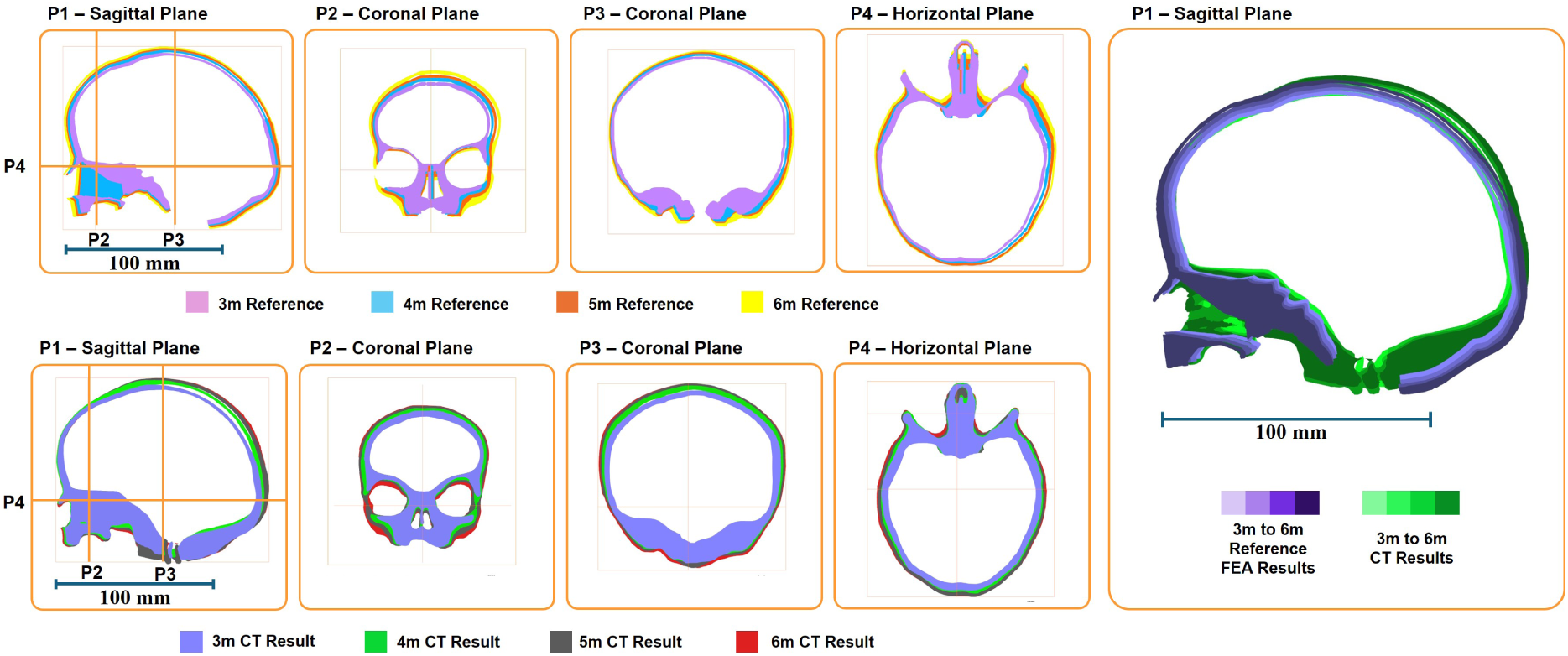
Two-dimensional sectional views from CT reconstructions comparing experimental and reference cranial morphologies. The figure includes: (a) CT-derived growth from 3–6 months; (b) reference growth from 3–4 months across four planes: P1 sagittal (midline profile), P2 coronal (anterior vault), P3 coronal (biparietal symmetry), and P4 horizontal (occipitofrontal extent); and (c) reference, CT, and overlapping images shown in three sagittal (P1) views.

## 4 Discussion

The observed trends in cranial morphometric measurements across all modalities show consistent incremental increases in BPD, OFD, and cranial heights (lateral: LCH; posterior: PCH; anterior: ACH), alongside ICV progression from approximately 619–695 mL at 3 months to 742–831 mL at 6 months. These patterns align with the expected postnatal cranial development driven primarily by encephalic expansion [40]. Manual measurements benefit from clear anatomical landmarks but remain vulnerable to operator variability as small differences in caliper placement can substantially affect results [39]. This subjectivity underscores the need for more precise and reproducible measurement techniques [52].

Multi-camera optical measurement using webcams offered an effective validation tool, generally consistent with manual and CT-derived values. However, discrepancies appeared in superior-view OFD and anterior-view ACH measurements. These were likely caused by vertical displacement of the skull during expansion, as internal water pressure induced both lateral widening and upward movement. This shift altered the pre-calibrated camera angles, obscuring or misaligning key landmarks. Consequently, multi-camera optical measurement underestimated cranial circumference due to the limited landmark visibility, and these values were not used (see Appendix D, Table 3). Despite these issues, errors remained systematic across ages, resulting in linear trends that still reflect overall growth. It should be noted that the camera-based system was employed primarily as an observational and secondary measurement tool, complementing rather than replacing the CT-derived volumetric data which served as the three-dimensional ground truth in this study. The inherent limitations of planar 2D imaging, including out-of-plane motion, perspective distortion, and parallax error, are well recognised for tracking a three-dimensionally expanding object. These limitations are reflected in the discrepancies observed in superior-view and anterior-view measurements and in the decision to measure cranial circumference manually rather than from camera footage. Where CT acquisition was available, it provided a direct three-dimensional measurement that is unaffected by these errors. The 2D camera system should therefore be understood in the context of this study as a real-time monitoring and supplementary surface measurement tool, whose known geometric limitations were mitigated by cross-referencing with CT-derived data.

Direct CT scanning has proven to be the most reliable method for cranial measurements. However, it poses challenges related to cost, access, and inability to visualize metallic features in future device developments.

ICV analysis showed linear regression trends with monthly increases of approximately 40–45 mL, indicating stable simulated growth. The slight deviation at 4 months arose because the reference ICV (754 mL) was taken from an individual healthy skull rather than the skull geometry from Liang et al. (743.13 mL) [1]. Beyond this exception, ICV values followed consistent, predictable growth trajectories.

A key advantage of the present dynamic platform over a set of discrete static age-specific phantoms is its ability to simulate the continuous process of cranial expansion within a single physical model. Rather than representing growth as a series of isolated snapshots, the system enables controlled volumetric actuation across the full developmental window from 3 to 6 months, allowing intermediate stages to be reproduced without fabricating additional models. Real-time monitoring of pressure, flow, and surface deformation during expansion provides data that a static phantom set cannot offer, including the trajectory of geometric change and the mechanical response of the phantom throughout the expansion process. Furthermore, producing a set of static phantoms for each age increment would require repeated fabrication, calibration, and assembly, introducing additional sources of variability and cost, factors which are particularly significant given that fabricating a single phantom currently requires approximately one month of production time. The present single-phantom approach therefore offers practical advantages in reproducibility, cost-effectiveness, and experimental efficiency, in addition to its capacity to capture the dynamic character of cranial development.

Growth patterns observed in manual anthropometry and multi-camera optical measurement showed both BPD and OFD increased steadily with age, consistent with normal cranial growth [5], driven by biological factors such as suture flexibility and intracranial pressure. In real pediatric skulls, vault enlargement is achieved through bone formation and remodelling at the suture margins. The present phantom instead reproduces this enlargement geometrically: a hyperelastic latex layer stretches under controlled internal pressure to reach each age-specific volume and dimension. A related structural limitation concerns the cranial bone plates themselves. In the fabricated phantom, the rigid resin plates can translate and reorient as the latex sutures stretch, but they cannot increase in size, since the model contains no mechanism for bone growth. In a real infant skull over this period, however, vault enlargement is achieved not only by expansion at the sutures but also by appositional bone growth and remodelling at the plate margins, so the plates themselves change in size and shape. The present phantom therefore accommodates the entire prescribed expansion through simulated suture (latex) stretching rather than through any growth of the bone plates. This simplification is appropriate for reproducing the overall geometry of normal vault expansion, but it is a particularly relevant limitation for conditions such as craniosynostosis, where one or more sutures fuse and growth is redistributed across the remaining plates and sutures. A fixed-plate phantom cannot capture this redistribution. The platform should therefore be understood as simulating the acute elastic deformation associated with cranial expansion rather than the underlying biological osteogenesis [58]. Water was used as the actuation medium to deliver controlled volumetric increments targeting age-specific ICV values; while this approach does not constitute a direct measurement or replication of encephalic pressure, it provides a reproducible and controllable loading condition for driving cranial expansion. Factors such as latex thinning during expansion, suture biomechanics, and physiological variations in internal pressure are not fully captured by this approach and represent acknowledged limitations of the current model. This distinction is particularly relevant for the device applications motivating this work: for some orthotic and protective devices, the intracranial pressure environment (not only the intracranial volume) is a critical loading parameter, and faithfully reproducing that pressure field would be a necessary step before the platform could be used to evaluate such pressure-dependent device behaviour. A further material consideration is that latex is a hyperelastic elastomer and does not reproduce the viscoelastic stress-relaxation and creep characteristic of fibrous biological cranial sutures. In principle, a viscoelastic suture held at a fixed deformation would exhibit gradual stress relaxation over time, requiring adjustment of the applied pressure to maintain a target intracranial volume. In the present study, however, the experimental protocol was designed to minimise the influence of this effect. Caliper-based measurements were completed within five minutes of reaching each target volume – a timescale over which stress-relaxation in latex is negligible. Camera-based measurements were captured instantaneously from continuous recordings. For CT acquisitions, the phantom was held at the target volume for approximately 50 minutes; however, the CT protocol required the phantom to complete a full rotation and return to its starting geometry, and no obvious image artefacts or reconstructed geometric distortion were observed. This provides indirect evidence that time-dependent deformation did not materially affect the phantom geometry during CT acquisition. Notwithstanding this, characterising the stress-relaxation and creep behaviour of the latex remains a limitation of the present material characterisation, and would be a necessary step for protocols involving prolonged pressure maintenance or cyclic loading.

Comparisons across modalities (Appendix D, Table 3) showed strong consistency, such as *<*1% variation in OFD, while highlighting differences such as multi-camera optical measurement overestimating BPD (e.g., 120.77 ± 0.29 mm vs. 118.71 ± 1.95 mm manual at 5 months), likely due to out-of-plane motion and parallax error. These findings, combined with manufacturing deviations shown in Fig. 3 (up to ∼3 mm) and growth-related distortions in Fig. 6 (0–4 mm), help characterize the reliability and limitations of each modality. Despite these minor imperfections, the synthetic skull growth model demonstrates strong potential for future research and methodological development.

The findings also have clinical relevance. By establishing a quantitatively validated baseline of normal cranial expansion – including trajectories of BPD, OFD, and ICV from 3 to 6 months – the platform provides a reference framework against which deviations associated with craniofacial conditions such as craniosynostosis could be contextualised in future work [34]. It should be emphasised, however, that both the reference data and the fabricated phantom represent normal cranial development; the platform has not been validated against pathological growth patterns such as those seen in craniosynostosis. As the therapeutic devices motivating this work are intended for atypical rather than normally developing skulls, a model explicitly validated to reproduce the relevant abnormality would ultimately be required for device evaluation in those conditions. The present normal-development platform therefore provides a validated baseline and a methodological foundation, rather than a disease-specific test-bed. From an anthropological perspective, the data highlight population-level variations in CI, such as mild brachycephalic trends from 81.8% to 87.7%, reinforcing the role of encephalic pressure in shaping early cranial morphology [59]. Beyond measurement, the controllable and CT-compatible nature of the platform makes it a promising test-bed for the future biomechanical evaluation of treatment options used in the management of craniosynostosis within this developmental window [30, 60, 61]. In its current form, the model reproduces the acute elastic response of the cranial vault to internal pressure rather than the active osteogenesis at the suture margins on which orthotic helmet therapy ultimately relies. Device-interaction studies on the present phantom should therefore be interpreted as characterising fit, contact, and short-term deformation under controlled loading, rather than the longer-term bone-remodelling response [31, 35]. Reproducing osteogenesis itself is a fundamentally biological process that lies beyond the scope of the present study, whose contribution is a validated, controllable, and CT-compatible mechatronic framework for replicating the geometry of cranial expansion.

Future work should extend the growth period beyond 6 months using a single physical phantom, enabling continuous simulation of cranial expansion over a longer developmental window. Improving phantom materials is another key direction, particularly materials that more closely mimic the nonlinear behavior of biological bone and sutures. It should be noted that the materials in the present phantom were selected to approximate the mechanical properties of the corresponding biological tissues as closely as practicable. However, application-specific material selection could further improve fidelity, but likely at the expense of the cost-effectiveness and accessibility that are central advantages of the current approach. Meta structure or architected-material designs may allow a degree of controlled widening of the bone surfaces under applied intracranial pressure, which would more closely approximate the mechanics of cranial expansion, though fully reproducing the underlying biological osteogenesis may remain beyond what an *in vitro* phantom can achieve. Advances in additive manufacturing could also be leveraged to fabricate more realistic, multi-material cranial models with spatially varying mechanical properties, further enhancing anatomical fidelity and enabling more sophisticated biomechanical studies. Limitations in manual and camera-based methods can be addressed using alternative three-dimensional surface measurement approaches, such as structured-light scanning, stereophotogramme-try, or markerless surface tracking systems. These would be particularly valuable in configurations where CT acquisition is unavailable or impractical, enabling full three-dimensional surface tracking without reliance on planar imaging.

This study demonstrated an anatomy-derived *in vitro* platform capable of replicating neonatal cranial growth from 3 to 6 months using a multi-material phantom, a precise pneumatic–hydraulic actuation system, and multi-modal measurement methods. The system produced consistent increases in cranial dimensions and intracranial volume, with strong agreement across manual, multi-camera optical, and CT-derived measurements, confirming its reliability for studying early cranial biomechanics. This platform provides a controllable and repeatable environment for evaluating growth patterns and supporting the future development and preliminary testing of clinical interventions. Future work will extend the growth period using a single phantom model and explore advanced materials and additive manufacturing approaches, including meta structures that better mimic biological bone and suture behavior, to create more realistic and adaptable cranial phantoms.

## Supporting information

Online Resource Video 1

## Appendices

### A Full Details of the Control Box and Graphical User Interface (GUI)

#### A.1 Control System (Pneumatic and Hydraulic Control Box)

The control box functions as the central hub for managing both pneumatic and hydraulic operations within the experimental setup. It enables precise regulation of pressure, flow, and actuation to simulate intracranial volume changes in the phantom skull model. The system integrates mechanical, electrical, and software components to provide accurate control and real-time monitoring, ensuring reproducible experiments that mimic physiological cranial expansion in infants.

A detailed schematic of the control box is shown in Fig. 7, illustrating the key components and their interconnections. Pneumatic lines are shown in black, hydraulic lines in light blue, electrical cables in blue, and signal lines in red (ground connections omitted for simplicity). The schematic presents the overall system architecture without exhaustive wiring detail.

**Fig. 7.**
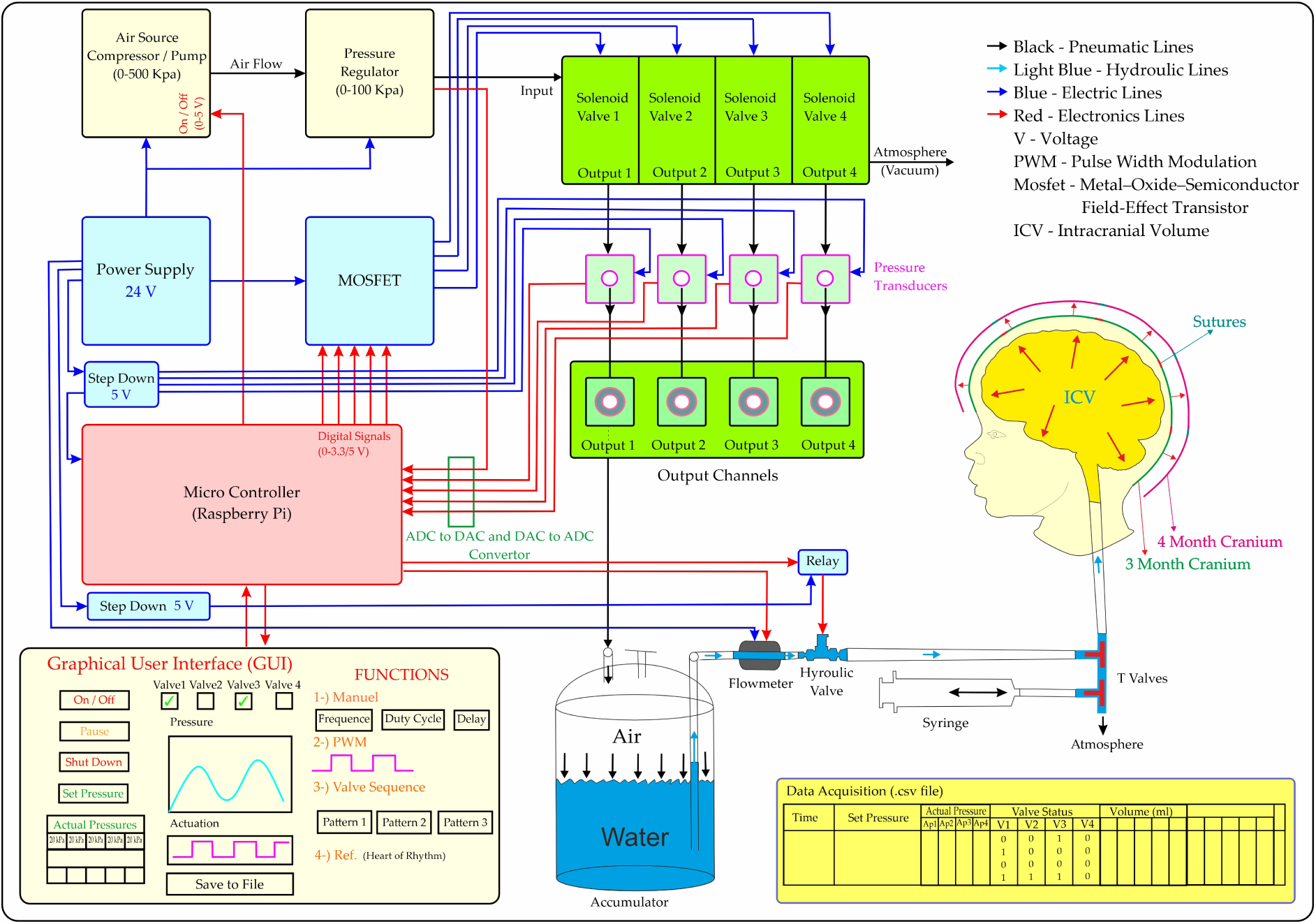
Detailed schematic of the control box components and connections.

**Fig. 8.**
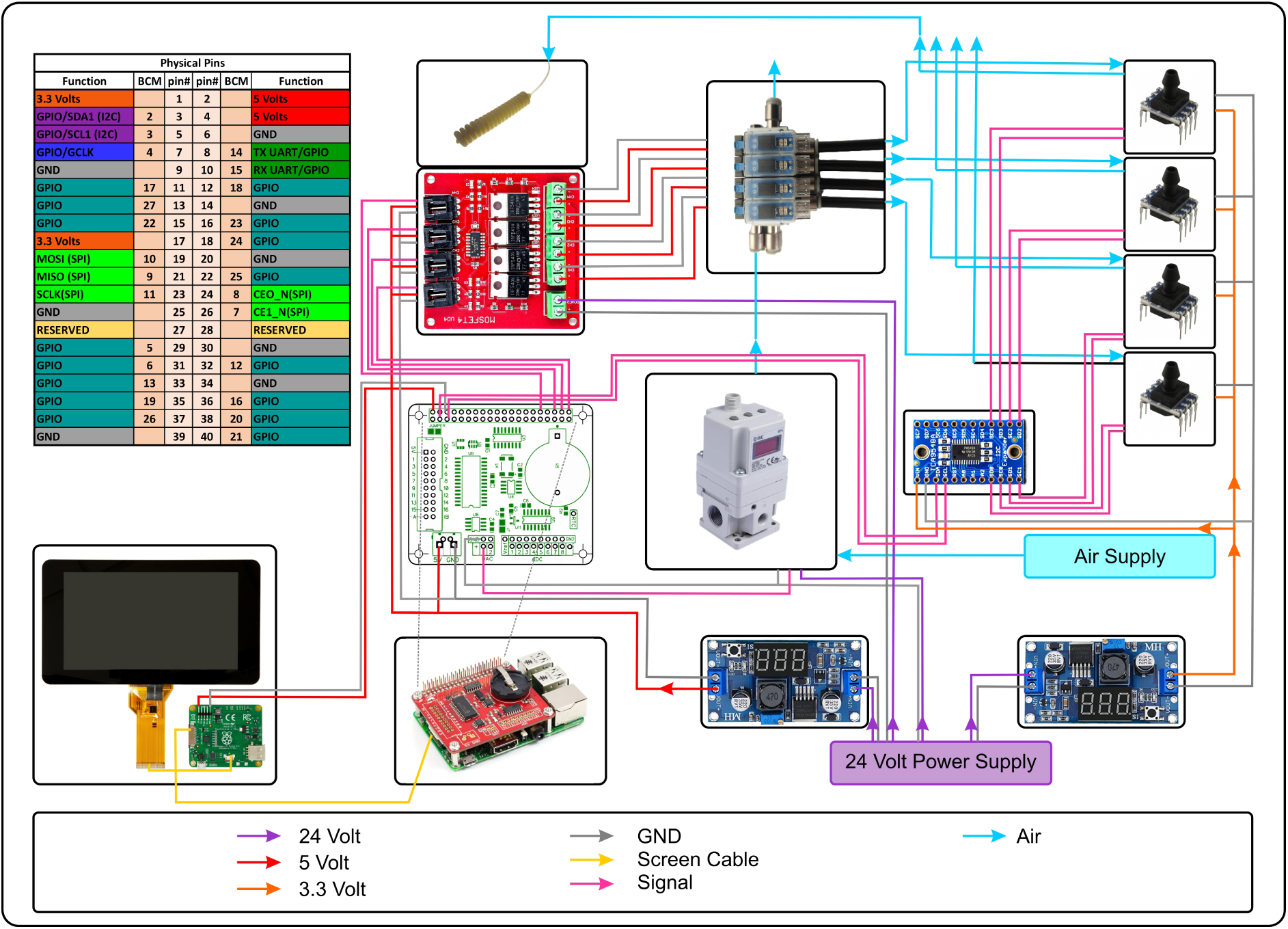
Pneumatic system design.

The system begins with an external compressed-air source or pump supplying constant airflow to a pressure regulator. The regulator adjusts the air to a defined pressure, which is then routed to the solenoid valves. A 24-V DC supply powers the system, with step-down regulators used to match individual component requirements. The pressure regulator and solenoid valves are controlled through a graphical user interface (GUI) running on a Raspberry Pi microcontroller. MOSFET transistors interface the micro-controller with the solenoid valves, while each valve connects to a pressure transducer via T-connectors for real-time pressure monitoring. Because the regulators and sensors operate using analog signals, an ADC/DAC module links the Raspberry Pi to these components.

Pressurized air from the solenoid valves fills an accumulator, which drives water through the tubing system. Before operation, the head model and tubing are evacuated using a syringe to remove air bubbles. Water is then introduced until the initial intracranial volume is reached (e.g., 694 mL for a 3-month-old), filling the head cavity without causing expansion. During this process, T-valves at the base of the phantom are briefly opened to vent small water amounts and ensure complete bubble removal. For the expansion phase (e.g., from 694 mL to 745 mL), the hydraulic valve admits a controlled volume of water while the corresponding pressure transducer provides continuous safety monitoring. The flowmeter supplies real-time volume feedback. When the target volume is reached, the hydraulic valve closes automatically via a relay triggered by the microcontroller. Throughout the procedure, pressure, flow, and actuation data are logged to a CSV file for post-experimental analysis.

#### A.2 Software and GUI Application

The control box software is implemented on a Raspberry Pi microcontroller and provides a user-friendly interface for system operation, parameter adjustment, and data visualization. The GUI – developed in Python using Tkinter or similar libraries – allows operators to set pressure thresholds, control valve states, and monitor system performance in real time.

The interface is organized into six numbered functional modules, corresponding to labels 1–6 in Fig. 9, in addition to a central actuation graph. The graph occupies the main window and plots valve-control voltage signals over time, using color-coded traces to distinguish channels. This enables real-time monitoring of both individual valve activity and coordinated multi-channel actuation.

**Fig. 9.**
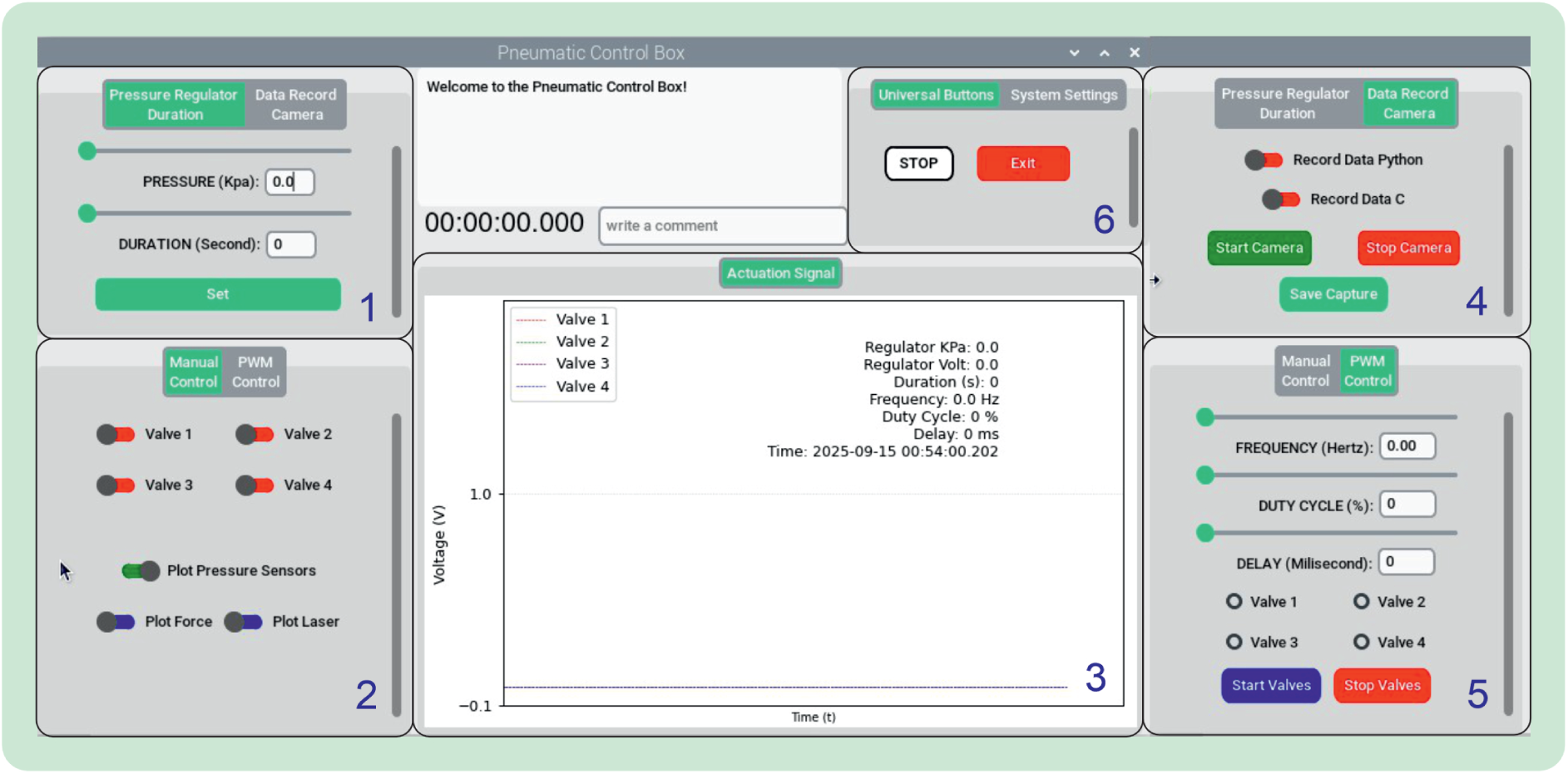
GUI for the pneumatic and hydraulic control box. Numbers 1–6 indicate the six functional modules described in Appendix A.2.

Module 1 allows configuration of the pneumatic pressure regulator and experiment duration. Pressures can be set between 0 and 100 kPa, and actuation time is defined in seconds, enabling precise specification of loading and timing parameters. Module 2 provides manual control of the four valves corresponding to the four pneumatic channels. It also offers options to view real-time data from pressure, force, and laser sensors, making it useful for calibration and manual testing. Module 3 displays the central actuation signal graph, showing real-time voltage responses of each valve channel throughout the experiment. Module 4 integrates data acquisition and video recording. Experimental data can be logged in Python or C formats, and synchronized video is recorded through a connected webcam. Start, stop, and save functions ensure both numerical and visual datasets are captured reliably. Module 5 supports automated actuation using pulse-width modulation (PWM). Users can specify frequency (Hz), duty cycle (%), and valve-to-valve delay (ms), enabling reproducible and complex actuation patterns. Module 6 provides system management functions. A universal stop button resets all active processes, and a safe-exit command closes the program to prevent unintended actuation and preserve data integrity.

### B Summary of Material Properties of the Latex, 3D Printed Components and Biological Tissues

The detailed mechanical properties of the materials used in the fabrication of the skull model are presented in the tables and figures below. The liquid dipping latex used for the production of the skin and sutures was manufactured at two different thicknesses and laser-cut into standardized specimens in accordance with ISO 37.

As illustrated in Fig. 10, uniaxial tensile tests were performed, and three specimens were tested for each thickness condition. During testing, time, displacement, and force data were recorded. Subsequently, the raw data were processed to obtain average stress–strain curves. The elastic modulus values were estimated from the initial low-strain region of the stress–strain curves using linear regression, corresponding to the initial tangent modulus, and are presented together with their standard deviations.

**Fig. 10.**
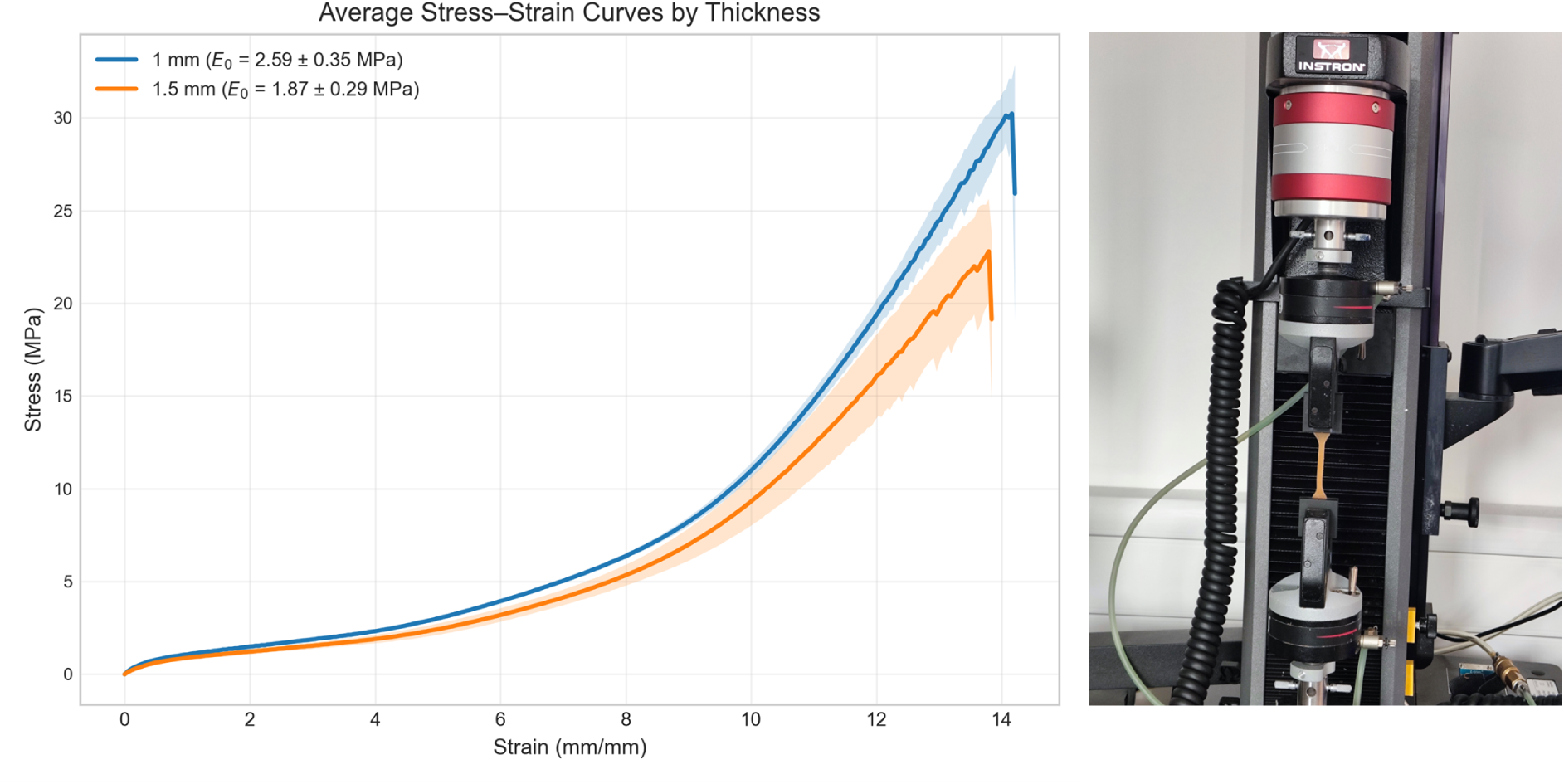
Uniaxial tensile testing setup and representative stress & strain curves for ISO 37 latex specimens of different thicknesses (1mm and 1.5mm).

**Fig. 11.**
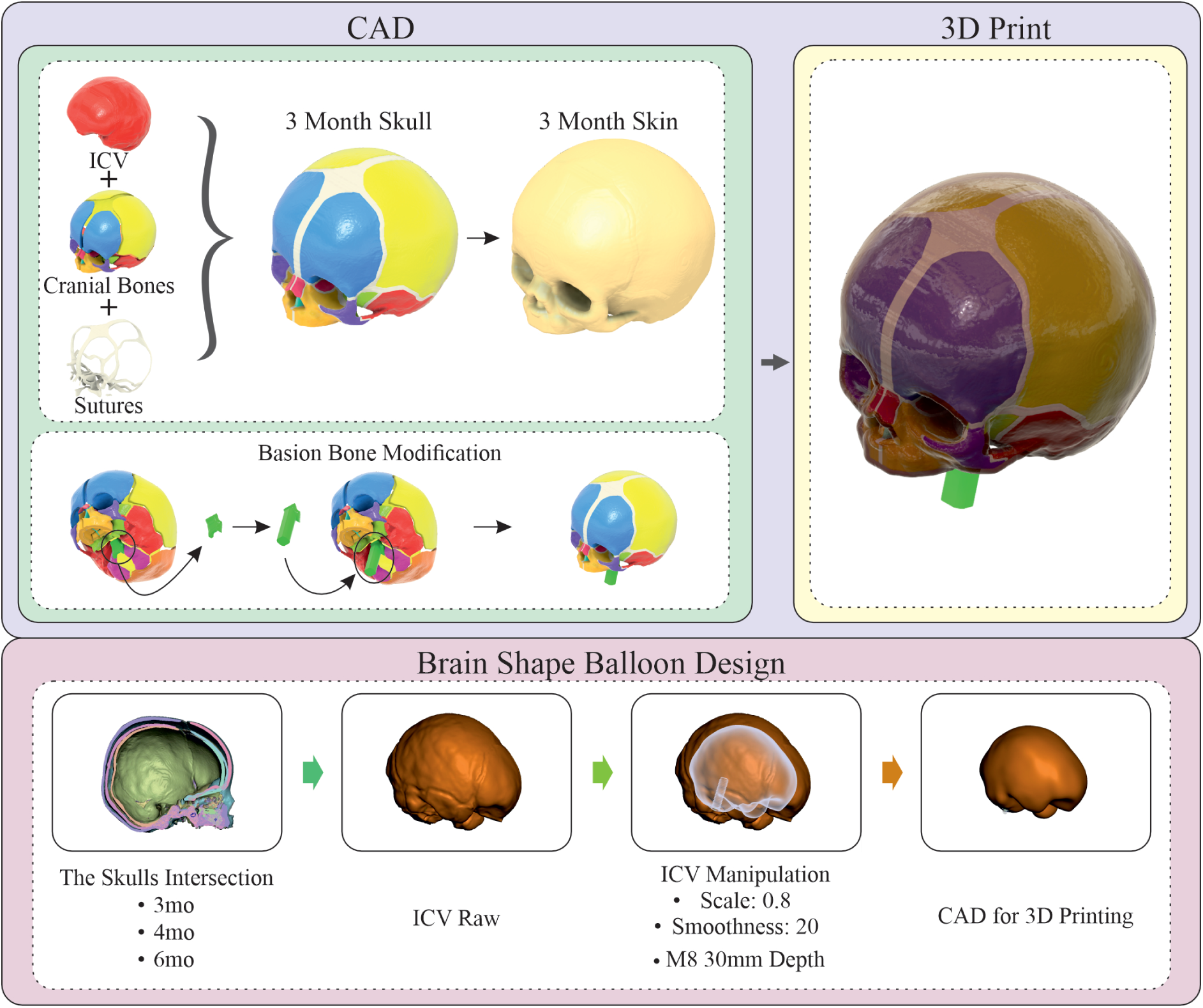
Fabrication workflow for the phantom head model and brain balloon. CAD integration of ICV, bones, and sutures produced the three-month skull with a 2-mm skin layer and basion fixation. The model was printed using Vero™ (bones) and Elastico™ (sutures/skin). The ICV geometry was scaled to 80% and smoothed to create the brain-shaped latex balloon via layer-by-layer casting.

Detailed results for each individual latex specimen, together with the corresponding averaged values and standard deviations, are presented in Table 1, which also includes the mechanical properties of the materials used for bone fabrication obtained from the official Formlabs material datasheets [41]. In addition, Table 2 provides a comparative overview of the mechanical properties of relevant biological tissues collected from the literature.

**Table 1.** Mechanical properties of latex (experimental, mean ± SD, *n* = 3) and resin materials (datasheet values).

| **Property / Materials** | **Latex Specimen (ISO 37, mean, 1 mm)** | **Latex Specimen (ISO 37, mean, 1.5 mm)** | **Tough 1500 Resin** | **Clear Resin V4.1** | **Grey Resin V4** |
|---|---|---|---|---|---|
| Thickness (mm) | 1 | 1.5 | – | – | – |
| Gauge Length (mm) | 30 | 30 | – | – | – |
| Width (mm) | 3 | 3 | – | – | – |
| Cross Section Area (mm<sup>2</sup>) | 3 | 4.5 | – | – | – |
| N Samples | 3 | 3 | – | – | – |
| Mean Initial Tangent Modulus (MPa) | 2.59 ± 0.35 | 1.87 ± 0.29 | – | – | – |
| Mean Max Stress (MPa) | 31.49 ± 1.01 | 25.50 ± 1.08 | – | – | – |
| Mean Max Strain (mm per mm) | 14.42 ± 0.23 | 14.72 ± 1.35 | – | – | – |
| Mean Max Force (N) | 94.47 ± 3.02 | 114.73 ± 4.84 | – | – | – |
| Mean Max Displacement (mm) | 432.72 ± 6.98 | 441.72 ± 40.64 | – | – | – |
| Elastic Modulus (MPa) | 2.59 ± 0.35 | 1.87 ± 0.29 | 1500 | 2300 | 2800 |
| Flexural Modulus (MPa) | – | – | 1400 | 2700 | 2200 |
| Ultimate Tensile Strength (MPa) | – | – | 33 | 53 | 65 |
| Elongation at Break (%) | – | – | 51 | 9 | 6 |

**Table 2.** Mechanical Properties of Biological Tissues from Literature.

| **Biological Tissue** | **Age Range** | **Elastic Modulus Range** |
|---|---|---|
| Bones | 3 Month – 6 Month | 186 – 3582 MPa |
| Sutures | 2 Day – 11 Month | 3.8 – 16.2 MPa |
| Skin | 2 Year – 94 Year | 20 – 25 MPa |

A wide range of values for the mechanical properties of biological tissues, particularly bone, have been reported in the literature [42, 62–66]. These variations arise due to multiple factors such as donor age, anatomical location of the extracted tissue, and testing conditions. In this study, material selection was primarily guided by age-specific data and mechanical properties relevant to cranial tissues. However, many studies lack detailed information regarding important factors such as moisture content, temperature, and the time elapsed between tissue extraction and testing. Furthermore, even for tissues obtained from the same anatomical region, significant variability in mechanical properties has been reported. Therefore, while literature does not allow for definitive conclusions, it provides a reasonable basis for approximate material matching.

Based on the data summarized in Table 2, the elastic modulus of skin samples obtained from donors aged between 2 and 94 years is typically reported in the range of 20–25 MPa [66]. Although this age range does not directly match the neonatal skull model used in this study, the maximum stress values of the latex samples (25.50–31.49 MPa) indicate that latex has the potential to mimic the mechanical response of skin.

For the sutures, the age range of the biological data reported in the literature (2 days to 11 months) is closely aligned with the developmental window modelled in this study (3 to 6 months) [67]. The mean initial tangent modulus of the latex (1.87–2.59 MPa) lies just below the range reported for biological cranial sutures (3.8–16.2 MPa) [67]. Two considerations qualify this comparison. First, the reported biological range is itself broad and highly condition-dependent: suture stiffness values in the literature vary not only with donor age but also with sex, specimen hydration and water loss, storage and preservation conditions for cadaveric tissue, temperature, the time elapsed between extraction and testing, and the testing protocol. The 3.8–16.2 MPa interval should therefore be regarded as an indicative estimate rather than a fixed physiological target. Second, the latex thickness in the suture regions of the fabricated skull is not uniform but varies as an inherent consequence of the manual dip-casting process. To quantify this, one representative skull was sectioned through the suture regions and the local latex thickness was measured with calipers, ranging from approximately 1 to 1.5 mm. Because the measured tangent modulus depends on thickness (2.59 MPa at 1 mm and 1.87 MPa at 1.5 mm; Table 1), the effective stiffness of the suture regions corresponds to this range of values rather than a single point. Taken together – and considering that the suture data correspond to the early end of the developmental window modelled here – the measured latex moduli can reasonably be regarded as a realistic approximation of neonatal suture behaviour at around three months of age, rather than as a poor match. The latex thus provides a compliant analogue of the suture: some-what softer than the mid-range literature values, but well within the uncertainty of the available biological data.

For bone tissue, the literature reports a wide range of elastic modulus values (186–3582 MPa) [32, 67]. The materials used in this study for bone fabrication—Tough 1500 Resin (1500 MPa), Clear Resin (2300 MPa), and Grey Resin (2800 MPa)—fall within this range [41]. In the final successful experiment, skull components were fabricated using Clear Resin.

Overall, these findings demonstrate that the selected materials provide a reasonable approximation of the mechanical properties of biological tissues, supporting the development of a representative skull model.

### C Full Details of the Initial Failed Fabrication Process of the Phantom Skull

#### C.1 Initial Design and Initial Manufacturing, Including Different Types of 3D-Printed Skulls and Problems

The development of the phantom skull model progressed through several iterative design and manufacturing stages aimed at achieving a realistic representation of infant cranial anatomy, with particular emphasis on cranial bones, intracranial volume (ICV) dynamics, and suture flexibility. Initial designs were created using computer-aided design (CAD) software and were informed by anatomical data extracted from medical imaging references to approximate the geometry of a pediatric skull.

The first CAD model integrated three core components: the ICV, the cranial bones, and the sutures. To approximate the external soft tissue envelope, a 2-mm offset skin layer was added around the skull geometry. Additionally, the basion bone was modified to function as a stable fixation point for the phantom, as this region exhibits minimal deformation during early cranial growth and therefore reduces unintended effects on experimental measurements.

The CAD skull models were fabricated using a Stratasys PolyJet 3D printer. A mul-timaterial printing strategy was adopted to approximate the heterogeneous properties of the pediatric skull: rigid Vero™ materials were used for the cranial bones, while Elastico™ photopolymer was selected for the sutures and skin layer to replicate compliant soft tissues. This combination allowed the printed phantom to better reproduce the mechanical behavior of cranial interfaces.

To create the intracranial component, the ICV geometry was adapted for the manufacture of a brain-shaped balloon. The raw ICV model was scaled to 80% of its original size and smoothed (factor = 20) to simplify manufacturing. The processed geometry was used to produce a latex balloon through a layer-by-layer casting process, providing a compliant and expandable structure capable of sustaining controlled intracranial pressurization.

Fig. 12 illustrates the assembly process. Model 1 shows the brain-shaped latex balloon. Model 2 depicts the modified 3D-printed skull, and Model 3 integrates the skull with a 2-mm offset skin layer. The balloon was incorporated into both configurations, enabling flexible testing of skull-only and skull-plus-skin models.

**Fig. 12.**
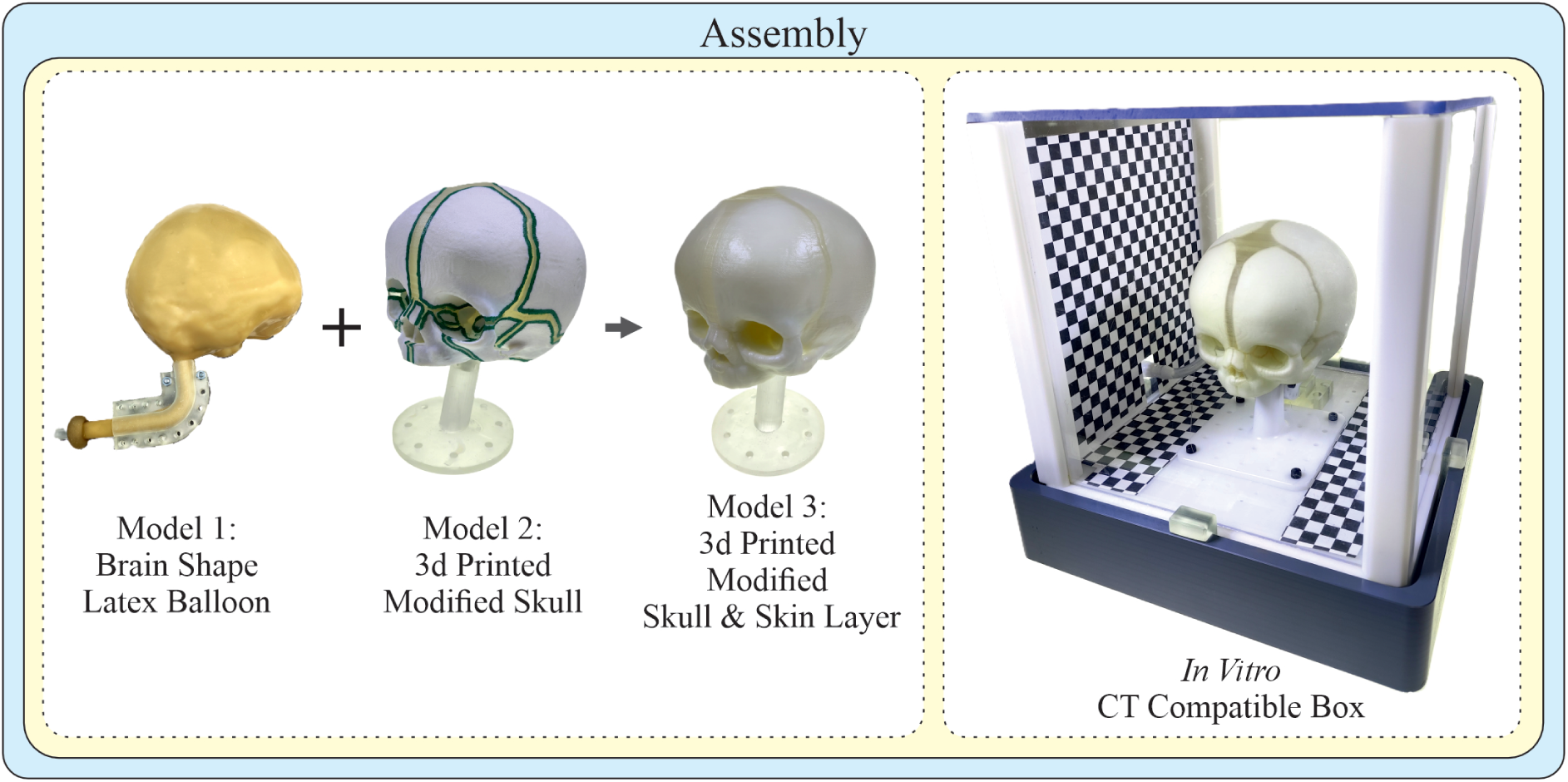
Assembly workflow of the phantom head model. Model 1 illustrates the latex brain balloon, Model 2 the modified 3D-printed skull, and Model 3 the skull with a 2-mm skin layer. Both configurations were mounted in a CT-compatible box with tubing for actuation, enabling stable and realistic intracranial expansion during initial tests.

Once assembled, the phantom was mounted inside a custom CT-compatible box that accommodated all tubing required for pneumatic actuation. This configuration enabled synchronized imaging and video capture while maintaining experimental stability. The assembled system performed effectively during initial trials, producing realistic cranial expansions and supporting reliable camera-based tracking without observable issues.

The experimental setup and an example of a failed phantom head model are shown in Fig. 13. During testing, one source of error was the entrapment of air between the surface of the brain-shaped balloon and the inner surface of the skull. Although this occasionally introduced measurement inaccuracies, it remained a minor issue. A more significant limitation arose from the mechanical performance of the Elastico™ material used for the sutures and skin. While initially flexible—allowing expansions equivalent to approximately one month of cranial growth—its elasticity deteriorated after two to three days. This degradation led to localized tearing in mechanically weak regions, particularly along the sutures. As the balloon expanded, pressure transferred to the cranial bones and concentrated at these vulnerable areas, ultimately causing irreversible deformation and model failure. To address this issue, Elastico™ was replaced with latex for the sutures and skin in later iterations, providing improved stretchability, durability, and long-term flexibility.

**Fig. 13.**
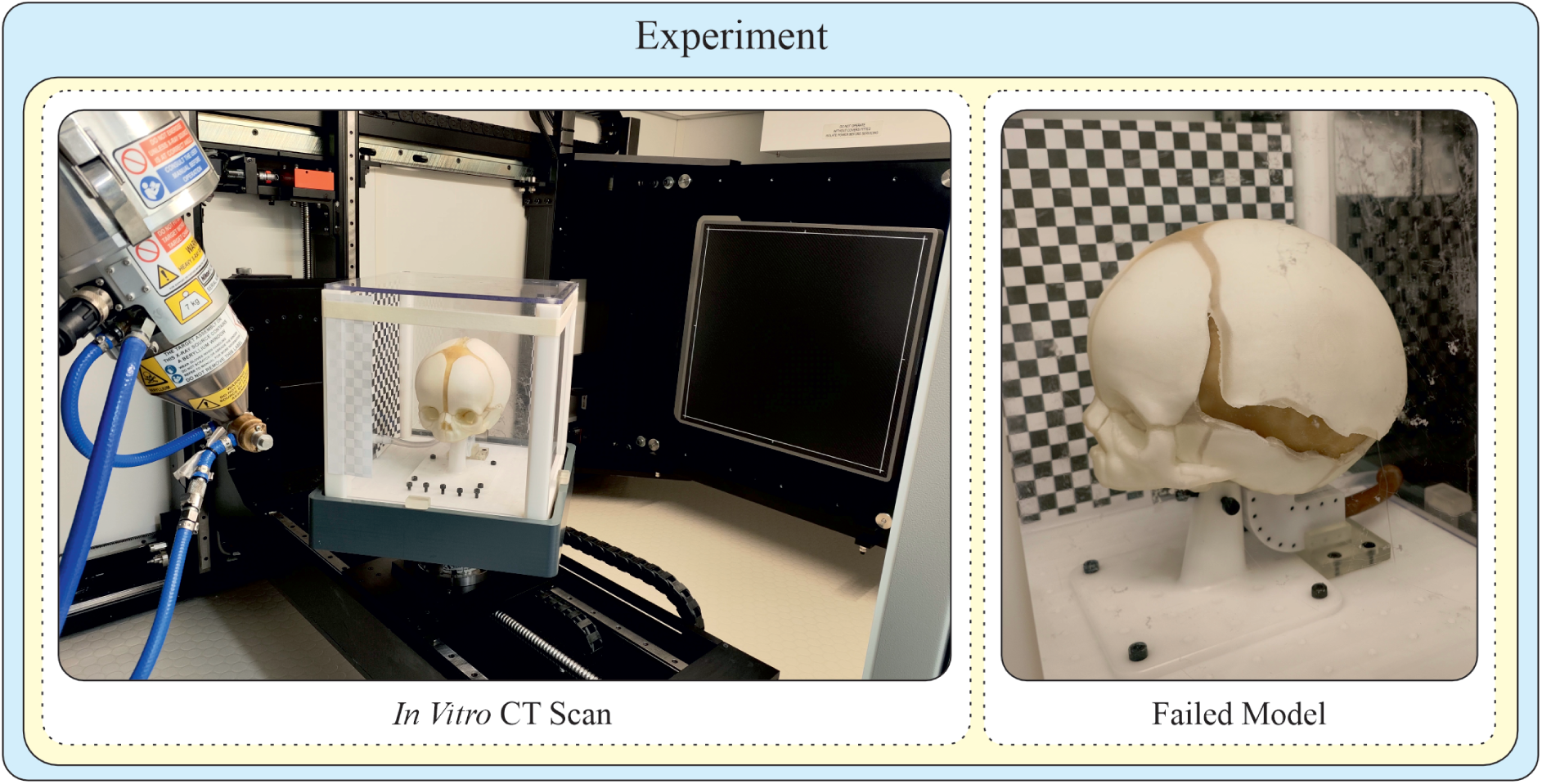
CT setup and example of a failed phantom. Left: phantom mounted in the CT-compatible box. Right: tearing in the suture regions caused by the loss of elasticity and stiffening of Elastico™ during expansion. Air pockets were occasionally present, but material degradation was the main failure mode. Later versions used latex for improved durability.

### D Summary of Cranial Morphometric Measurements

The summary of cranial morphometric measurements obtained using different methods is presented in Table 3. Manual and optical values are mean ± SD of three repeated measurements (*n* = 3), whereas CT values represent a single acquisition per age. At the 3-month baseline, BPD and OFD served as the pixel-to-millimetre scale references for each camera view and are therefore identical to their CT values with zero SD, as is the cephalic index derived from them. Cranial width is listed under the superior, posterior, and anterior views, and cranial length under both the superior and lateral views; where a modality acquires a dimension once, the same value appears in each view.

**Table 3.** Summary of cranial morphometric measurements. Manual and optical values are mean ± SD (*n* = 3); CT values are from a single acquisition per age. In the column headers, green and orange correspond to the transverse and vertical measurement lines used for the camera views in Fig. 4. In the manual rows, each linear dimension was measured once and then listed under every view in which it appears: lilac cells are the biparietal diameter and rose cells the occipitofrontal diameter. In the optical rows, blue cells are the CT-derived values used for pixel-to-millimetre calibration at the 3-month baseline (SD = 0 by construction), and cream cells indicate measurements considered unreliable owing to parallax and out-of-plane displacement of the skull during expansion. BPD = biparietal diameter; OFD = occipitofrontal diameter; LCH = lateral cranial height; PCH = posterior cranial height; ACH = anterior cranial height; CC = cranial circumference; CI = cephalic index. All linear values are in mm; CI in %.

| Measurement Method | Age | Superior View |  | Lateral View |  | Posterior View |  | Anterior View |  | Global Measures |  |
|---|---|---|---|---|---|---|---|---|---|---|---|
|  |  | BPD | OFD | OFD | LCH | BPD | PCH | BPD | ACH | CC | CI |
| Manual cranial anthropometry (mean ± SD, *n* = 3) | 3 months | 113.35 ± 0.86 | 137.13 ± 1.81 | 137.13 ± 1.81 | 104.59 ± 0.99 | 113.35 ± 0.86 | 111.45 ± 0.65 | 113.35 ± 0.86 | 104.21 ± 0.61 | 400.33 ± 2.52 | 82.67 ± 0.53 |
|  | 4 months | 116.30 ± 1.62 | 139.24 ± 1.60 | 139.24 ± 1.60 | 106.63 ± 1.30 | 116.30 ± 1.62 | 115.60 ± 1.50 | 116.30 ± 1.62 | 106.39 ± 0.86 | 407.00 ± 4.00 | 83.53 ± 0.77 |
|  | 5 months | 118.71 ± 1.95 | 141.65 ± 1.47 | 141.65 ± 1.47 | 106.88 ± 1.02 | 118.71 ± 1.95 | 117.14 ± 0.33 | 118.71 ± 1.95 | 107.41 ± 1.06 | 414.00 ± 5.00 | 83.80 ± 0.71 |
|  | 6 months | 120.32 ± 0.85 | 143.09 ± 1.27 | 143.09 ± 1.27 | 108.88 ± 1.80 | 120.32 ± 0.85 | 120.47 ± 1.75 | 120.32 ± 0.85 | 109.34 ± 0.33 | 416.67 ± 4.04 | 84.09 ± 0.24 |
| Multi-camera optical measurement (mean ± SD, *n* = 3) | 3 months | 113.14 ± 0.00 | 129.12 ± 0.25 | 138.29 ± 0.00 | 103.39 ± 0.30 | 113.14 ± 0.00 | 112.03 ± 0.26 | 113.14 ± 0.00 | 128.25 ± 0.30 | 381.91 ± 0.35 | 81.81 ± 0.00 |
|  | 4 months | 116.41 ± 0.52 | 131.62 ± 0.47 | 139.84 ± 0.34 | 105.98 ± 0.36 | 117.32 ± 0.34 | 115.13 ± 0.45 | 114.20 ± 0.54 | 127.00 ± 0.54 | 391.42 ± 0.54 | 83.24 ± 0.26 |
|  | 5 months | 120.77 ± 0.29 | 133.32 ± 0.53 | 140.84 ± 0.67 | 108.15 ± 0.32 | 120.52 ± 0.70 | 118.65 ± 0.23 | 115.10 ± 0.82 | 124.92 ± 1.00 | 401.52 ± 0.54 | 85.76 ± 0.34 |
|  | 6 months | 124.31 ± 0.56 | 135.19 ± 0.70 | 141.68 ± 0.69 | 109.88 ± 0.65 | 124.24 ± 0.46 | 121.87 ± 0.82 | 115.98 ± 0.30 | 121.87 ± 0.85 | 409.38 ± 0.78 | 87.74 ± 0.12 |
| Computed tomography (CT, single acquisition) | 3 months | 113.14 | 138.29 | 138.29 | 104.46 | 113.14 | 112.63 | 113.14 | 104.67 | – | 81.81 |
|  | 4 months | 116.37 | 140.54 | 140.54 | 106.15 | 116.37 | 115.70 | 116.37 | 107.13 | – | 82.80 |
|  | 5 months | 118.58 | 142.02 | 142.02 | 108.06 | 118.58 | 118.75 | 118.58 | 109.04 | – | 83.50 |
|  | 6 months | 120.01 | 143.45 | 143.45 | 109.36 | 120.01 | 120.50 | 120.01 | 110.54 | – | 83.66 |

## Acknowledgements

The authors would like to thank Ahmet Muhammet Kocabıyık for his assistance in debugging and refining the software used in the pneumatic and hydraulic control box.

## Author Contributions

All authors contributed to the conception and design of the study. Material preparation, experimental setup development, and analysis were performed by Ahmet Arif Colakoglu. The first draft of the manuscript was written by Ahmet Arif Colakoglu, and all authors commented on previous versions of the manuscript. All authors read and approved the final manuscript.

## Declarations

### Data Availability

The datasets generated and analysed during the current study are available from the corresponding author upon reasonable request.

### Ethics Approval

This study did not involve human participants, human tissue, or live animals.

### Consent for Publication

All authors consented to the publication of this manuscript.

